# Bimodal stimulation for the reduction of tinnitus using vibration on the skin

**DOI:** 10.1101/2022.11.28.518290

**Authors:** Michael V. Perrotta, Izzy Kohler, David M. Eagleman

## Abstract

Tinnitus (ringing in the ears) affects 1 in 10 adults in the United States, often with damaging psychological consequences. Currently, there exists no cure for most forms of tinnitus. Recently, bimodal stimulation – the pairing of sounds with haptic stimulation – has shown efficacy in reducing the symptoms of tinnitus. Previous bimodal stimulation approaches have used electrical shocks on the tongue, a technique that requires daily in-person sessions at an audiologist’s office. We here show that excellent results can be achieved wearing a wristband with multiple vibratory motors. Tones are played and the wristband correspondingly vibrates the wrist of the user at different spatial locations depending on the frequency of the tone. We compared the experimental group with a control group who listened to the tones but did not wear the wristband. The tone frequencies were centered on each user’s tinnitus frequency and the tones were randomized both in frequency and duration. 45 participants with Tinnitus Functional Index (TFI) scores of 25 and above were tested. Results show a significantly greater reduction in TFI scores for the experimental group compared to the control. Importantly, with higher baseline severity we find larger differences between the experimental and control groups, revealing greater symptom improvement for those with severe tinnitus. The therapeutic approach of combining sounds with spatially- and temporally-correlated vibrations on the wrist is found to be a simple, time-efficient, and effective procedure to lessen the symptoms of tinnitus.

## Introduction

Tinnitus is the perception of sounds without the presence of a corresponding sound source (Langguth et al., 2019). Individuals who experience tinnitus commonly describe it as constant and irritating sounds in the ears that can make it difficult to concentrate, to understand what people are saying, and to hear external sounds (Langguth et al., 2019; Tang et al., 2019). Tinnitus can have various etiologies, many of which are closely tied to neurological insult and hearing loss; common examples include sensorineural hearing loss (Kaltenbach, 2011), noise induced hearing loss (Kaltenbach et al., 2000), presbycusis (Kang et al., 2021), ototoxicity from oral medications (Cianfrone et al., 2011), neurologic and vestibular disorders, and infectious disease (Langguth et al., 2019). Tinnitus affects 21.4 million adults (9.6%) in the United States, a third of whom experience near-constant symptoms (Bhatt et al., 2016). It occurs in over 1 in 4 Americans between the ages of 65 and 85 (KochKin et al., 2011). Tinnitus is correlated with a variety of comorbidities, including hypertension and depression (Park et al., 2022). Tinnitus can be experienced in different ways (ringing, static, pulsatile, etc) and at varying levels of severity. For 10% of patients, tinnitus severity is enough to elicit relentless frustration, annoyance, anxiety, depression, cognitive dysfunction, insomnia, stress, and emotional exhaustion (Langguth et al., 2019).

Generally, models suggest tinnitus is a result of the brain’s attempt at compensating for a loss of sensory auditory information (De Ridder et al., 2011). No treatment currently exists to completely remove tinnitus. Treatments which help alleviate the symptoms of tinnitus include counseling (e.g. cognitive behavioral therapy), pharmacological treatments (e.g. lidocaine), hearing aids, sound masking, and brain stimulation (e.g. transcranial magnetic stimulation) (Langguth et al., 2013).

Bimodal stimulation is a relatively new treatment used to alleviate the symptoms of tinnitus. Researchers demonstrated that pairing auditory tones with small shocks on the tongue reduced the symptoms of tinnitus (Conlon et al., 2020, 2022; Marks et al., 2018; Riffle et al., 2021). In those studies, the tongue was chosen based on the hypothesis that bimodal stimulation requires synchronous activity in the dorsal cochlear nucleus, where information from the ears first converges with haptic stimulation from the head and neck (Kaltenbach et al., 2005); therefore, tactile stimulation of the tongue seemed to be the appropriate haptic input. Next, other researchers demonstrated that bimodal stimulation works with a variety of timing characteristics between shock and tone and that the treatment had high compliance and satisfaction rates (Conlon et al., 2020, 2022), although it should be noted that this paper did not present a control group.

We set out to determine whether tones combined with haptic stimulation – not on the tongue but on the skin of the wrist – would work just as well. If so, this would challenge the hypothesis that the bimodal signals need to converge at the dorsal cochlear nucleus, and would provide an easier treatment modality for the reduction of tinnitus.

## Methods

We recruited adults in the United States via an online advertisement for a study investigating a new tinnitus treatment. Interested participants completed the Tinnitus Functional Index questionnaire (TFI) (Henry et al., 2016; Meikle et al., 2012) and were included if they scored 25 points or higher. A total of 45 participants were randomly assigned to the experimental condition (bimodal stimulation with tones + wristband) or a control condition (tones only). The research protocol was approved by the IRB and written consents were obtained from all participants.

Each participant completed ten minutes of daily treatment over the course of the eight-week study. The treatment included listening to tones and, unless in the control condition, feeling corresponding vibrations on the wrist. All participants completed the TFI questionnaire each week during the eight-week period.

Tones were played through a phone app in a random order and with random timing. Each tone had a duration between 150 ms and 350 ms with an inter-tone silence of 0 ms to 500 ms. Each tone played without replacement before repeating the sequence in a newly shuffled order. The range of tone frequencies was set to be specific to the participant’s main tinnitus frequency, which was collected in the app before the start of the study. Their tinnitus frequency was calibrated as the center of their individualized tone range. During treatment, 120 tones per minute played at frequencies spread logarithmically between an octave above and below each participant’s customized tinnitus frequency.

Treatment was administered through a phone app (sounds) and wristband (vibrations on the skin). Participants opened the app and listened to tones for ten minutes each day. Experimental participants listened to tones while feeling corresponding vibrations on their wristband (Figure 1). The app played the tones while sending spatially- and temporally-corresponding vibrations to the wristband via Bluetooth.These vibrations were spread evenly across the wrist in a logarithmic manner, such that an octave spanned the same spatial distance regardless of its frequency. Although the wristband has only four vibrating motors, frequencies were mapped onto one of 256 different spatial locations, accomplished by leveraging a haptic illusion (Alles, 1970; Luzhnica et al., 2017; Rahal et al., 2009). Specifically, an illusory location is a point on the wrist that is felt by the wearer of the wristband even when a motor is not located directly at that point. We stimulate these illusion locations by turning on two motors, one on either side of the illusion location, at specific amplitudes such that the wearer feels as if a single point somewhere between the two motors is vibrating.

**Figure 1.**
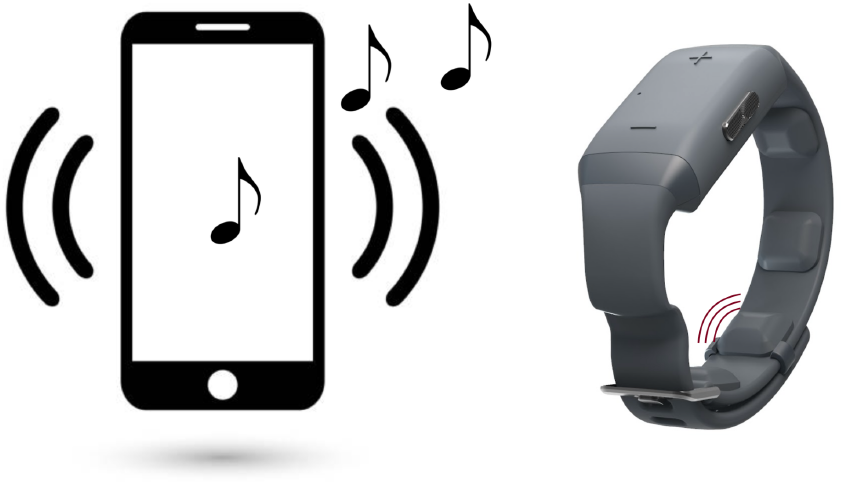
Bimodal stimulation. Tones are played from a phone app; a haptic wristband vibrates simultaneously with each tone. The location of the vibration is a function of the frequency of each individual tone, such that the frequency map is spread across the skin. Participants in the control condition listened to tones from the phone app but did not receive a wristband and were given no haptic stimulation.

The audio-only control participants did not receive a wristband and only listened to the tones without feeling any vibration.

## Results

Participants in the experimental group (tones paired with simultaneous vibrations of the wristband) showed a clinically-significant average improvement in TFI scores of -17.9 (SD = 18.17, n = 22, p<.001, two-tailed t-test; **Figure 2A**). Interestingly, the audio-only control group also showed a significant difference from baseline of -7.5 (SD = 15.35; n = 23; p=.03, two-tailed t-test); however, it is important to note that a change in TFI score of less than -13 is not considered clinically significant (Meikle et al., 2012), so while the audio-only control showed a change, it was not clinically meaningful. The difference between the experimental and control groups was statistically significant (t(43) = 2.10, p=.04).

**Figure 2.**
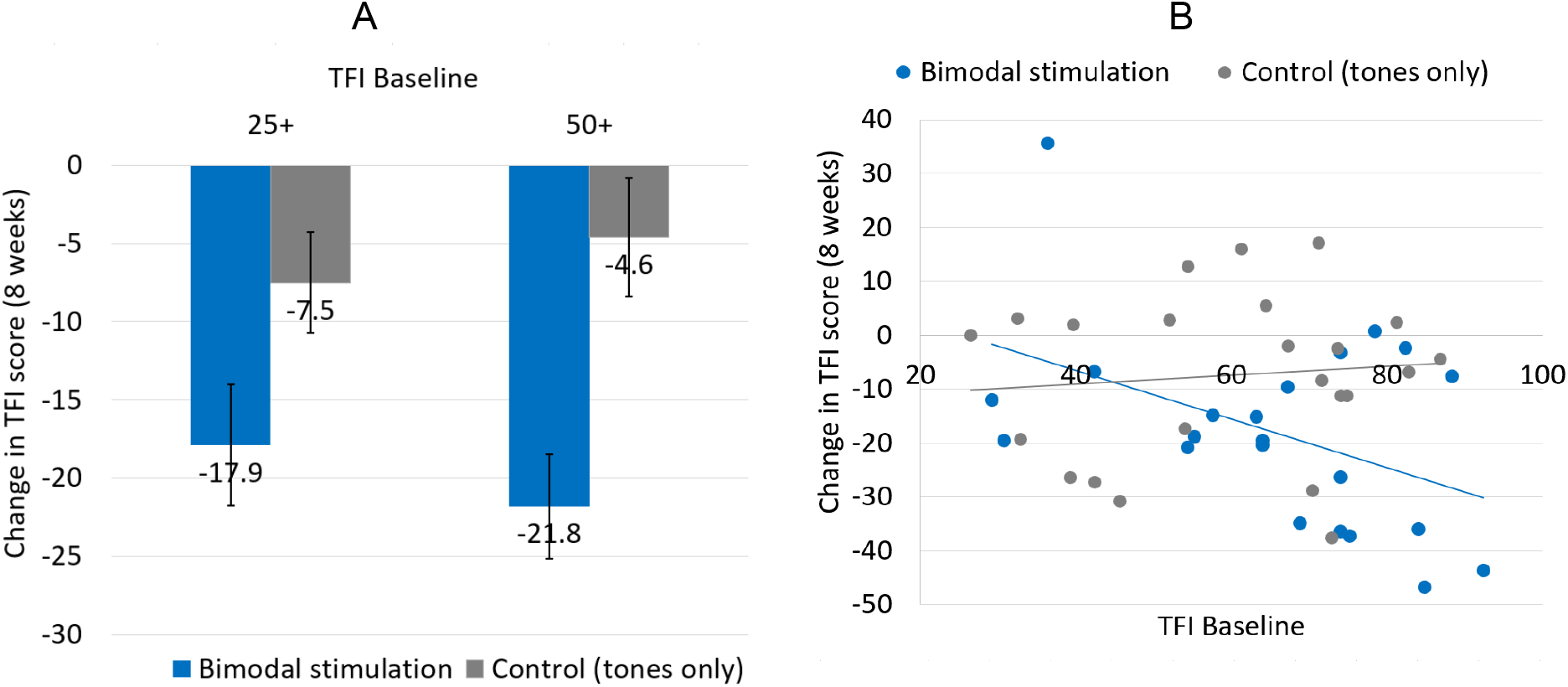
Bimodal stimulation with the wristband lowers tinnitus symptoms, especially for those with upper-moderate to severe tinnitus. **(A)** The average change in TFI score after eight weeks of treatment was significantly greater for experimental participants when compared to tone-only control participants (left; p=.04). When performing this comparison only on participants with TFI scores greater than or equal to 50, the difference is greater and more statistically significant (right; p=.002). Error bars show SEM. **(B)** Change in TFI score after 8 weeks of treatment plotted against baseline TFI score. For the experimental group, a higher baseline TFI score predicts a larger improvement. The same correlation was not present in the control group.

Importantly, participants who started the study with greater severity of tinnitus (as determined by their baseline TFI score) experienced greater improvement from the bimodal stimulation. The difference between the experimental and control groups increased with higher baseline TFI scores: for baselines of 50 and above, the experimental group averaged -21.8 (SD=14.6; n=18; p<.001) while the control group averaged -4.6 (SD=15.0; n=18; p=.24); the significance of the difference increased to p=0.002 (two-tailed, t(32) = 3.39).

(Meikle et al., 2012) defined tinnitus as a “significant problem” in the TFI range from 25 to 50 and “severe” for any score above 50 points. In a more recent analysis of the TFI score distribution, “lower moderate” tinnitus was between 18 and 42 points, “upper moderate” was between 42 and 65 points, and “high” is any score higher than 65 points (Gos et al., 2021).

The correlation of a larger reduction with a higher TFI baseline can be seen by plotting improvement of each participant against baseline TFI score (**Figure 2B**). This effect is not present with the control group The weekly progression of participant scores can be seen for all participants in **Figure 3**. For all participants in the experimental group, 91% showed improvements (i.e. TFI moved in the negative direction), and 64% of those improvements qualified as clinically significant (TFI score reduction of 13 points or greater; (Meikle et al., 2012) (**Figure 3A**). When restricting analysis to those subjects with moderate/severe tinnitus (baseline TFI 50 and above), 94% of these participants showed improvements (in the negative direction), and 72% of those improvements qualified as clinically significant (**Figure 3C**). In contrast, in the control group, 52% showed improvement and 30% of those qualified as clinically significant (**Figure 3B**); when restricting to those with baseline TFI of 50 and above, 63% showed improvements and 19% of those improvements qualified as clinically significant (**Figure 3D**)

**Figure 3.**
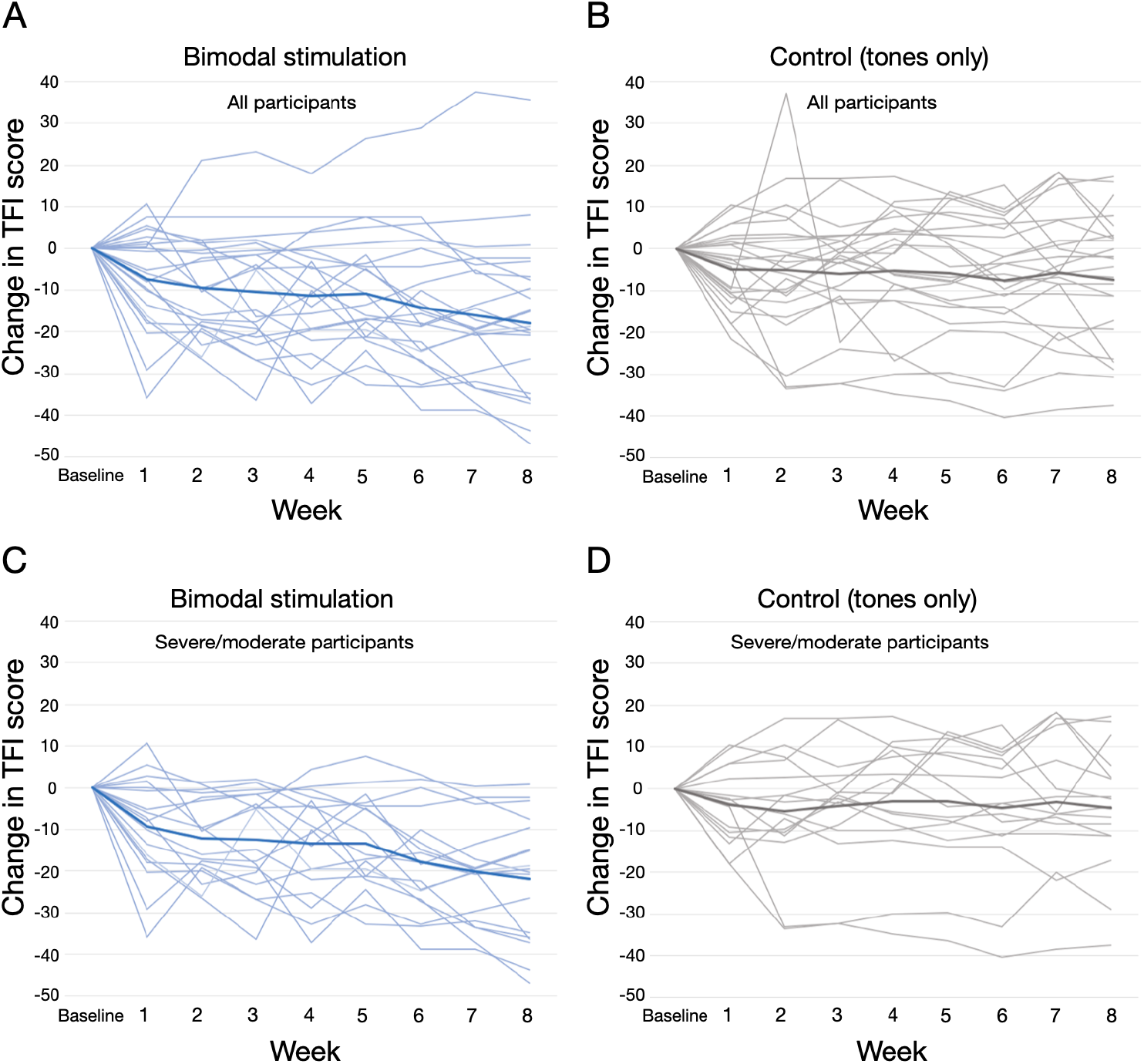
Changes from TFI baseline over 8 weeks for upper-moderate and severe tinnitus. (A) Experimental group (tones + vibrations) with all participants included (TFI baseline 25+). (B) Control group (tones only) with all participants included (TFI baseline 25+). (C) Experimental group for participants with TFI baseline 50 and above. (D) Control group for participants with TFI baseline 50 and above. Individual participants are shown as thin lines and the average as thick lines.

We note that many participants in both the experimental and control groups had some amount of variance in their TFI scores from week to week. Presumably, this is at least partially a function of the broad range of tinnitus etiologies. Although we searched, we found no significant correlations between self-reported tinnitus etiology (or any other collected demographic data such as age, duration of tinnitus, or hearing loss) and TFI score change.

## Discussion

We found improved TFI scores for both the experimental (tones and vibration) and control (tones only) groups. However, only the experimental group’s average TFI improvement was clinically significant (greater than 13 points).

Importantly, individuals with a greater severity of tinnitus at baseline experience a larger improvement in their tinnitus with bimodal treatment (**Figures 2 and 3**). For example, participants starting at a baseline TFI score of 50 and above demonstrate an almost five-fold greater improvement than those treated with tones only (**Figure 2A**).

It is worth noting the size of the effect in the audio-only control experiment. It has long been known that there is a placebo effect in tinnitus treatments (Duckert & Rees, 1984), and our control-group results may be an expression of that. However, despite the effect of tones alone, the average result was well below clinical significance, and the result was highly significantly smaller than tones-plus-haptics.

The experimental group showed improvements in their TFI score that match or exceed previous reports employing electric shocks on the tongue instead of vibrations on the wrist. Specifically, (Conlon et al., 2020) found TFI improvements at week 12 between -10 and -15 points for three experimental groups, while (Marks et al., 2018) found an average TFI improvement of -8 points over 4 weeks. In comparison, our experimental group showed improvement of -17.9 for all participants, and -21.8 for those with moderate to severe tinnitus (baseline TFI score 50+).

Among other things, these results suggest that convergence of neural signals at the dorsal cochlear nucleus is not the site of action of the improvement, as the same or better average TFI improvement is found with tactile stimulation of the wrist. Thus, the current findings open the door to an easier and less expensive method of tinnitus treatment.

The results of this study lay the framework for future research to investigate how different patterns and frequency ranges of tones affect the efficacy of treatment. Conlon et al (2022) showed that changing the parameter settings after 6 weeks of treatment allowed study participants to continue to make progress for an additional 6 weeks. This was an improvement on the findings of their prior study in which study participants experienced the greatest amount of progress in the first 6 weeks, but then plateaued for the final 6 weeks. Their study ended at 12 weeks, but potentially changing parameter settings at regular intervals could allow continued progress to resolution.

## Conclusion

We have demonstrated that bimodal stimulation using audio tones (individualized to a participant’s tinnitus frequency) combined with spatially-corresponding vibrations on the wrist can significantly reduce the symptoms of tinnitus more than a control treatment of tones alone. Those with more severe tinnitus at baseline experience larger improvements, almost five times that of the control group.

